# Melanoma cells suppress mast cell growth via a melanin-dependent mechanism

**DOI:** 10.1101/2025.02.03.636204

**Authors:** Fabio Rabelo Melo, Lea Nyman, Ida Österman Menander, Mirjana Grujic, Gunnar Pejler

## Abstract

Mast cells (MCs) have a well-established detrimental role in allergic conditions, but they can also impact on diverse malignant conditions, including melanoma. To study the latter, previous studies have mainly evaluated how MCs can influence melanomas/melanoma cells. However, the inverse scenario, i.e., whether melanoma/melanoma cells might impact on MCs has received less attention. Here we investigated this issue and show that melanoma cell-conditioned medium had a strong growth-inhibitory impact on MCs, which was attributed to inhibition of MC proliferation combined with induction of apoptosis. Further, our data indicate that such effects were attributable to melanin present in the melanoma cell-conditioned medium, as similar anti-proliferative effects were seen in response to both free melanin and to melanocores enriched from melanoma-conditioned medium. Melanin did not reduce the expression of MC markers, but was shown to impair MC activation. We also demonstrate that melanin is taken up by MCs, both in cultured MCs and *in vivo* in melanoma tumors, and it was observed that melanin, after uptake, can be found in the MC nucleus. Further, we show that melanin had marked effects on the nuclear morphology in MCs accompanied by clipping of core histone 3, and it is demonstrated that these events were dependent on translocation of tryptase, a granule-localized protease, into the MC nucleus. Tryptase was also shown to affect the mechanism of melanin-induced cell death. Altogether, the present study outlines a novel mechanism by which melanoma cells can suppress MC function, potentially representing an immunosuppressive mechanism that may influence tumor growth.

## Introduction

Mast cells (MCs) are immune cells characterized by a remarkably high content of secretory granules, containing various pro-inflammatory compounds such as histamine, growth factors, cytokines, proteases and serglycin proteoglycans ^1^. When MCs are activated, e.g., through IgE receptor crosslinking, these compounds are released to the exterior and can cause a powerful inflammatory reaction ^2, 3, 4^. Such responses can be further amplified through *de novo* production of additional pro-inflammatory compounds following MC activation ^3^.

MCs can contribute to our host defense against parasites and bacterial infection, and they also have an important role in the clearance of various toxins ^5, 6, 7, 8^. However, MCs are also known to have detrimental functions in various pathological contexts, most notably exemplified by allergic settings, where MCs represent major effector cells ^2, 3, 4^. In addition, MCs have been implicated in malignant disease. In support of the latter, MCs are known to infiltrate various types of tumors, including breast cancer, prostate cancer, lung cancer, lymphomas and also melanoma ^9, 10, 11, 12^. In malignant settings, MCs predominantly localize to the tumor stroma, but they can also be found within the tumor parenchyma ^9, 10, 11, 12^.

Considering the abundance of MCs in tumor tissues, it is reasonable to assume that MCs may have a functional impact on malignant processes. Indeed, there is now substantial documentation supporting such a notion, both from clinical and experimental studies ^9, 10, 11, 12^. For melanoma, there are several clinical studies showing that MCs are abundant in the tumor tissue, and it has also been reported that MC density shows a correlation with tumor prognosis. In several such cases, MC presence has been associated with poor outcome ^13, 14, 15, 16, 17, 18^ whereas other studies have linked MC presence to a good prognosis ^19, 20^. In animal studies, some reports have suggested that MCs can support melanoma growth ^21, 22^ whereas others suggest the opposite, i.e., that MCs may offer protection against melanoma progression^23, 24, 25^.

As to the mechanism by which MCs can influence melanoma, it has been suggested that MCs can promote tumor growth by secreting pro-angiogenic factors such as VEGF ^16, 26, 27^, FGF-2 ^15^ and histamine ^26, 28^, and it has also been shown that MC presence correlates with vessel density in melanomas ^14, 15, 16, 29^. Further, it has been reported that MCs can support melanoma growth through their expression of TIM-3 ^30^, HIF-1a ^26^ or complement factor 3 ^31^. Other reports suggest that MCs may augment melanoma growth by promoting myeloid-derived suppressor cell activity ^32^ or by enhancing the resistance to anti-PD-1 immunotherapy ^33^ ^34^. In contrast to these studies in which various MC products/mechanisms are suggested to support melanoma growth, there are also reports revealing that MC products can have a protective function in melanoma. For example, it has been shown that MC tryptase can have a dampening effect on melanoma cell proliferation, as seen both in experimental studies ^24, 35^ and in clinical studies in which tryptase levels correlated with favorable melanoma prognosis ^36, 37, 38^. Further, it has been demonstrated that MCs can suppress melanoma metastasis by inhibiting HMGA1 secretion ^22^.

As outlined above, there is a now wealth of evidence linking MCs to melanoma, and substantial efforts have been undertaken to investigate how MCs impact on melanoma/melanoma cells. However, it is largely unknown as to whether there is a bidirectional interaction between melanoma cells and MCs, i.e., whether melanoma cells can have an influence on MCs. For example, an outstanding question is whether melanomas can exert immunosuppression by impacting on tumor-associated MC populations. Here we addressed these issues. Our findings reveal that melanoma cells have a profound growth-reducing impact on MCs, and that this effect is largely attributable to melanin secreted by the melanoma cells. Potentially, this could represent a novel principle for immunosuppression in melanoma.

## Materials and Methods

### Animals

Mice (8-18 weeks old; males/females) deficient in mMCP-6 ^39^ and wild-type (WT) controls were on C57BL/6J genetic background. Animal experiments were approved by the local ethical committee.

### B16.F10 cell line

B16.F10 mouse melanoma cells were cultured in Dulbecco’s modified Eagle’s medium (Gibco), supplemented with 10% heat-inactivated fetal bovine serum (Invitrogen), 50 μg/ml streptomycin sulfate (Sigma-Aldrich), 60 μg/ml penicillin G (Sigma-Aldrich), 2 mM L-glutamine (SVA, Uppsala, Sweden). Cells were kept at 37°C in 5% CO_2_ and medium replaced when cells achieved 90 to 100% confluency.

### Bone marrow-derived MCs

Femurs and tibiae from mice of same gender and age were removed and MCs were obtained by culturing bone marrow cells in Dulbecco’s modified Eagle’s medium (Gibco), supplemented with 10% heat-inactivated fetal bovine serum (Invitrogen), 50 μg/ml streptomycin sulfate (Sigma-Aldrich), 60 μg/ml penicillin G (Sigma-Aldrich), 2 mM L-glutamine (SVA) and 10 ng/ml mouse recombinant IL-3 and SCF (Peprotech). Cells were kept at 0.5 x 10^6^ cells/ml at 37°C in 5% CO_2_ and the medium was replaced every 3 to 4 days. Cells were used after 4 weeks of culture, and were shown to represent pure MC populations as based on morphological criteria and toluidine blue staining. MCs were cultured under normal conditions (Control) or in medium containing 30% B16.F10-conditioned medium (Cond. Medium). Control experiments showed that MCs cultured in 70% normal MC medium/30% of medium used for melanoma cell culture (see above) show similar growth rate and similar viability as cells grown in 100% MC medium.

### EdU labeling

0.5 x 10^6^ cells/ml were distributed individually into a 24-well plate. Cells were treated with 30% B16.F10 melanoma-conditioned medium or different concentrations of melanin/melanocores and kept at 37°C for 48h. Four to 12h before harvesting the cells, 10 μM of EdU was added. Cells were then harvested by centrifugation (400 x g, 5 min) and stained using the Click-iT^TM^ Plus EdU Alexa 647 flow cytometry kit (ThermoFisher), followed by flow cytometry analysis (BD Accuri C6 plus, BD). Data from 10,000 events/sample were collected and analyzed by the FlowJo software (BD biosciences).

### Cell viability

Cells were harvested, washed once in PBS, resuspended in Annexin binding buffer (BD Pharmingen), stained with AnnexinV-FITC and Draq7 and analyzed by flow cytometer (BD Accuri C6 plus, BD).

### Cell Morphology and counts

Transmission Electron Microscopy (TEM) was performed as previously described ^40^, and for light microscopy assays cells were collected onto cytospin slides and stained with a Fontana-Masson staining kit (Sigma-Aldrich) in combination with toluidine blue (Sigma-Aldrich) or DAPI (ThermoFisher). Images were acquired with a Nikon 90i microscope equipped with DIC and NIS-Elements software (Nikon). Ten µl aliquots of cell suspensions were mixed with the same volume of trypan blue solution. Cell size and counts were performed by using the Countess II FL cell counter (Life Technologies).

### Histology of tumor sections

B16.F10 melanoma tumor sections from paraffin-embedded tissue were deparaffinized and hydrated. Sections were stained with Fontana-Masson in combination with toluidine blue for 5 min, followed by washing with distilled water. Images were acquired by using a Nikon Ni-U microscope equipped with NIS-Elements software (Nikon).

### Melanocore enrichment

Melanocores from B16.F10 conditioned medium were obtained by filtration using the centrifugal filter units Amicon Ultra-15 ultracel-100k (Millipore). Tubes (15 ml) were centrifugated (4000 x g; 25°C; 20 min).

### Western blot analysis

Equal number of cells were collected and solubilized in Laemmli buffer (Bio-Rad). Thereafter, samples were subjected to SDS-PAGE using Bio-Rad Mini-Protean TGX stain-free gels, followed by transfer to 0.2 μm nitrocellulose membrane (Trans-Blot turbo transfer pack – Bio-Rad) using the Bio-Rad Trans-Blot Turbo transfer system. Membranes were then blocked with Intercept blocking buffer (Li-Cor), incubated with primary antibody/antiserum overnight (4°C), followed by incubation with IRDye secondary antibody (Li-Cor) for 1h at room temperature. Membranes were scanned by using an Odyssey CLx scanner (Li-Cor).

### Cell fractionation

MCs (1 x 10^6^ cells) were collected by centrifugation, washed with PBS and subcellular protein fractions were prepared by using the Subcellular Protein Fractionation Kit for Cultured Cells (Thermo Scientific) according to manufacturer’s specifications. All samples were resuspended in 100 μl of Laemmli buffer (Bio-Rad) and analyzed by Western blot.

### Endocytosis inhibition

One ml of MC suspensions (0.5 x 10^6^ cells/ml) were distributed in triplicates into a 24-well plate and samples were either left untreated or treated with 50 μg/ml of melanin or melanocores, in the presence or absence of 3 μM MitMAB or 10 μM Dyngo-4a. Endocytosis inhibitors were added 1h prior the melanin/melanocore treatment. Cell viability was assessed after 24h using Annexin V-FITC and Draq7 staining/flow cytometry (BD Accuri C6 plus, BD). Cytospin slides were prepared and stained with Fontana-Masson and DAPI. Images were acquired by using a Nikon 90i fluorescent microscope equipped with DIC filter.

### Quantification of melanin

Melanin concentration was measured by using a Melanin Assay Kit (Sigma-Aldrich) according to the manufacturer’s instructions. Alternatively, melanin content was measured by using a protocol where melanoma cell-conditioned medium was monitored for absorbance at 335 nm, and by comparison with a standard curve constructed by adding defined amounts of melanin to non-conditioned medium.

### Peritoneal cell -derived mast cells (PCMCs)

Mice were euthanized, and the abdominal skin was carefully removed. Seven milliliters of sterile Tyrode’s buffer were injected into the abdominal cavity, and peritoneal cells were collected using an 18G needle syringe. The cells were rinsed twice with Tyrode’s buffer and once with complete medium (DEM; Gibco) supplemented with 10% heat-inactivated fetal bovine serum (Invitrogen), 1% MEM (Sigma-Aldrich), 50 μg/ml streptomycin sulfate (Sigma-Aldrich), 60 μg/ml penicillin G (Sigma-Aldrich), 2 mM L-glutamine (SVA), 50 μM β-mercaptoethanol (Gibco) and 20 ng/ml mouse recombinant IL-3 and SCF (Peprotech). Peritoneal cells from two mice were cultured separately at 0.5 x 10^6^ cells/ml, and after three days, the cultures were combined. The cells were maintained at 0.5 x 10⁶ cells/ml at 37°C in 5% CO₂, with the medium replaced every 3 to 4 days.

### Tunel assay for in situ apoptosis detection

Paraffin-embedded B16.F10 melanoma tumor sections ^24^ were stained using the Click-iTTM Plus TUNEL Assay Kit with Alexa 488 (Invitrogen) for in situ apoptosis detection, following the experimental protocol for tissue sections. After TUNEL staining, the slides were washed twice with PBS containing 1% BSA and blocked with 10% goat serum in PBS for 30 minutes at room temperature. The slides were then washed twice with PBS/1% BSA and incubated overnight at 4°C with 100 μl of rabbit anti-Mcpt6 immune serum (1:500) in PBS/1% BSA, with or without pre-immune serum at the same concentration. Next, the slides were washed three times with PBS and incubated with a goat anti-rabbit Alexa-555-conjugated antibody in PBS/1% BSA for 1 hour at room temperature, protected from light. The slides were then washed three times with PBS/1% BSA and stained with Hoechst 33342 (NucBlueTM, Invitrogen) for 10 minutes, followed by three washes with PBS. Finally, the slides were mounted using SlowFade® Gold Antifade Mounting Medium (Life Technologies) and analyzed with a Nikon Eclipse 90i upright fluorescence microscope equipped with Nikon NIS-Elements software.

### Mast cell activation

One ml of MC suspensions (0.5 x 10^6^ cells/ml) were distributed in triplicates into 24-well plates and samples were either left untreated or treated with 50 μg/ml of melanin or melanocores. For IgE-stimulated degranulation, IgE anti-DNP (0.1μg/ml) was added alone or together with melanin or melanocores, followed by overnight incubation. DNP-HSA (0.5 μg/ml) was added 2 h before the end of the 24 h experiment. Compound 48/80 activation was performed by adding 50 μg/ml of compound 48/80 2 h before the end of the experiment. Samples were collected by centrifugation (400 x g, 5 min), washed in PBS/1% BSA and pellets were resuspended in 300 μl of the same buffer. Anti-mouse CD63-PE-Cy7 antibody and/or isotype control (Invitrogen) were added and incubated for 1h at 4°C. Cells were then washed in PBS/1% BSA and resuspended in 300 μl of the same buffer, followed by flow cytometry analysis (BD Accuri C6 plus, BD biosciences). Data were analyzed using FlowJo software (BD biosciences).

### Laser-scanning microscopy

100 µl aliquots from 10^6^ cells/ml suspensions were dropped into round areas on microscopic glasses prepared with liquid-repellent slide marker pen. Suspensions were settled for 15 min and liquid was removed with filter paper. Cells were dried for 15 min and fixed with 4% paraformaldehyde in PBS (15 min), followed by a 15-min drying step. 100 μl of 0.1% Triton-X100 in PBS was added to each individual glass and incubated for 10 min at room temperature. Next, 100 μl of rabbit anti-Mcpt6 immune serum (1:500) in TBS/1% BSA and/or pre-immune serum at the same concentration were added and incubated overnight at 4°C, followed by washing 3 times with TBS. Anti-rabbit Alexa-488-conjugated antibody in TBS/1% BSA was added and incubated for 1h at room temperature. The slides were kept in dark, washed 3 times with TBS. Finally, slides were stained with Hoechst 33342 (NucBlue^TM^, Invitrogen) for 10 min, followed by 3 times washing with TBS. Slides were mounted with SlowFade® gold antifade mounting medium (Life Technologies). For assessment of nuclear structure, slides were prepared as described above, fixed with 4% paraformaldehyde, permeabilized and stained with 0.5% saponin solution containing 1 μM TO-PRO-3 nuclear fluorescent dye. Samples were analyzed using a LSM 710 laser-scanning microscope equipped with ZEN 2009 software (Carl Zeiss). Z-stack sections were acquired and used for 3D image assembly and analyzes by using Imaris software (Oxford Instruments).

### Quantitative real time RT-PCR

Total RNA was extracted by using NucleoSpin RNA kit (Machery-Nagel), following the manufacturer’s protocol. cDNA was prepared by using the iScript cDNA synthesis kit (Bio-Rad) and stored at -20°C. RT-PCR was performed by using the iTaq universal SYBR green supermix kit (Bio-Rad) and analyzed in triplicates on 384-well microplates (Bio-Rad) using the CFX384 Touch RT-PCR machine (Bio-Rad). Gene expression was calculated by using the software CFX Maestro and expressed as relative to Gapdh (Bio-Rad). The following primers were used: Gapdh, forward: 5′- CTCCCACTCTTCCAC CTTCG-3′; Gapdh, reverse: 5′-CCACCACCCTGTTGCTGTAG-3′; Mcpt6, forward: 5′-CATTGATAATGACGAGCCTCTCC-3′; Mcpt6, reverse: 5′- CATCTCCCGTGTAGAGGCCAG-3′; CPA3, forward: 5′- TGACAGGGAGAAGGTATTCCG-3′; CPA3, reverse: 5′-CCAAGGTTGACTGGATGGTCT-3′; Serglycin, forward: 5′-GCAAGGTTATCCTGCTCGGAG-3′; Serglycin, reverse: 5′- GGTCAAACTGTGGTCCCTTCTC-3′; TNFα, forward: 5′- CCCTCACACTCAGATCATCTTCT-3′; TNFα, reverse: 5′-GCTACGACGTGGGCTACAG-3′; VEGFa, forward: 5′-GCACATAGAGAGAATGAGCTTCC-3′; VEGFa, reverse: 5′- CTCCGCTCTGAACAAGGCT-3′; FGF2, forward: 5′- GCACATAGAGAGAATGAGCTTCC-3′; FGF2, reverse: 5′-CTCCGCTCTGAACAAGGCT-3′; IL6, forward: 5′-AGACAAAGCCAGAGTCCTTCAGAGA-3′; IL6, reverse: 5′- TAGCCACTCCTTCTGTGACTCCAGC-3′; TGFβ, forward: 5′- CTCCCGTGGCTTCTAGTGC-3′; TGFβ, reverse: 5′-GCCTTAGTTTGGACAGGATCTG-3′

## Results

### Melanoma-conditioned medium inhibits MC proliferation

To address the possible impact of melanoma cells on MCs, we first prepared conditioned medium from B16.F10 melanoma cells and then cultured a low number (0.1 x 10^6^ cells) of bone marrow-derived MCs with or without such conditioned medium (30% B16.F10-conditioned medium mixed with normal culture medium) continuously for 7 days (without replacing the cell culture medium, which was possible due to the low number of cells). As seen in Fig. 1A, MCs grown without the addition of melanoma-conditioned medium expanded markedly during the culture period, showing an approximately 7-fold increase in numbers after the 7-day period. In contrast, MCs grown in the presence of melanoma-conditioned medium did not display any detectable cell growth.

**Fig. 1.**
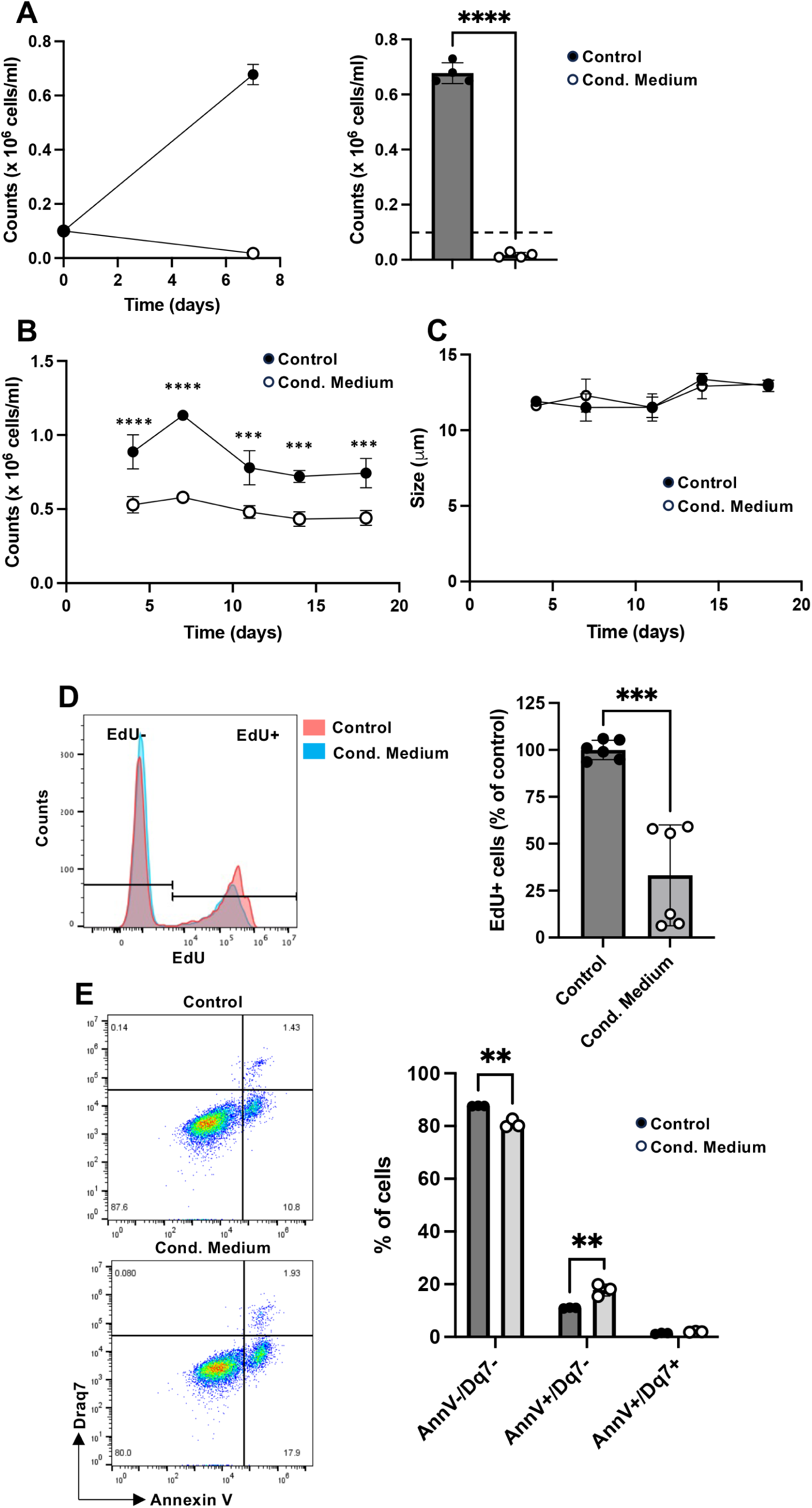
Melanoma-conditioned medium inhibits MC proliferation. MCs were cultured under normal conditions (Control) or in medium containing 30% B16.F10-conditioned medium (Cond. Medium). (A) MCs were cultured for 7 days, and cell counts were performed at day 0 and after 7 days in culture; the right panel shows quantification of cell counts after 7 days. (B) MC cultures were followed for 18 days. Medium was replaced every 3^rd^/4^th^ day, cells were counted and diluted to 0.5 x 10^6^ cells/ml after each passage. (C) Cell size in μm. (D) Cell proliferation assay based on EdU incorporation. (E) Cell viability assessment using Annexin V (AnnV) and Draq7 (Dq7) staining. Viable cells (AnnV^-^/Dq7^-^), apoptotic cells (AnnV^+^/Dq7^-^) and late apoptotic/necrotic cells (AnnV^+^/Dq7^+^). The data are representative of at least three independent experiments and are given as mean values ± SEM (n=3). Unpaired t-test; \**p*≤0.05; \*\**p*≤0.01; \*\*\**p*≤0.005; \*\*\*\**p*≤0.0001.

We next used an approach in which MCs were seeded at a higher density (0.5 x 10^6^ cells/ml). Cells were then cultured for 3 days, after which cells were recovered, washed, quantified and again seeded out at 0.5 x 10^6^ cells/ml (after dilution, if applicable) in fresh culture medium. The same procedure was then repeated every third/fourth day. These analyses revealed that MCs cultured in the absence of melanoma-conditioned medium at each assessed time point showed a clear expansion in cell numbers, typically showing a ∼2-fold increase in numbers (i.e., from 0.5 x 10^6^ cells/ml at the starting point to ∼1 x 10^6^ cells/ml at the end of each 3/4rday period)(Fig. 1B). In contrast, when MCs were grown in the presence of melanoma-conditioned medium, no detectable expansion of the cell populations could be seen at any assessed time point (Fig. 1B). Instead, cell numbers remained at a constant level. It was also noted that the culture of MCs in the presence of melanoma-conditioned medium did not affect the size of the MCs (Fig. 1C).

To address the mechanism underlying the blockade of MC growth by melanoma-conditioned medium, we assessed whether this could be explained by effects on proliferation. Indeed, as shown by EdU staining, melanoma-conditioned medium caused a profound reduction of MC proliferation (Fig. 1D). We next asked whether the reduction in MC growth additionally could be linked to an increased population of non-viable cells, and this was assessed by Annexin V/Draq7 staining. This analysis revealed that the melanoma-conditioned medium caused a slight, yet significant increase in the proportion of non-viable MCs, as reflected by an increase in the population of Annexin V^+^/Draq7^-^ (apoptotic) cell population (Fig. 1E). Notably, no significant increase in Annexin V^+^/Draq7^+^ (necrotic) cells was observed in response to melanoma-conditioned medium. These data thus suggest that melanoma-conditioned medium suppresses MC growth by combined inhibition of proliferation and induction of apoptosis.

### Melanin and melanocores impair MC growth through antiproliferative and pro-apoptotic mechanisms

We next aimed to elucidate the mechanism by which the melanoma-conditioned medium affects MC growth. Melanin is a main component of melanoma-conditioned medium, and we thus considered the possibility that the inhibitory action of melanoma-conditioned medium on MCs might be attributable to its content of melanin. To address this, we estimated the concentration of melanin in the melanoma-conditioned medium, and then added corresponding amounts of purified melanin to the MCs. These analyses indicated that the melanin content was ∼10-50 µg/ml in the used mixture of B16.F10-conditioned medium and normal medium, and when MCs were cultured in the presence of melanin at 50 µg/ml, cell growth was profoundly decreased (Fig. 2A). Significant inhibition of cell growth was seen starting from 10 µg/ml, and at 100 µg/ml the effect of melanin was further enhanced.

**Fig. 2.**
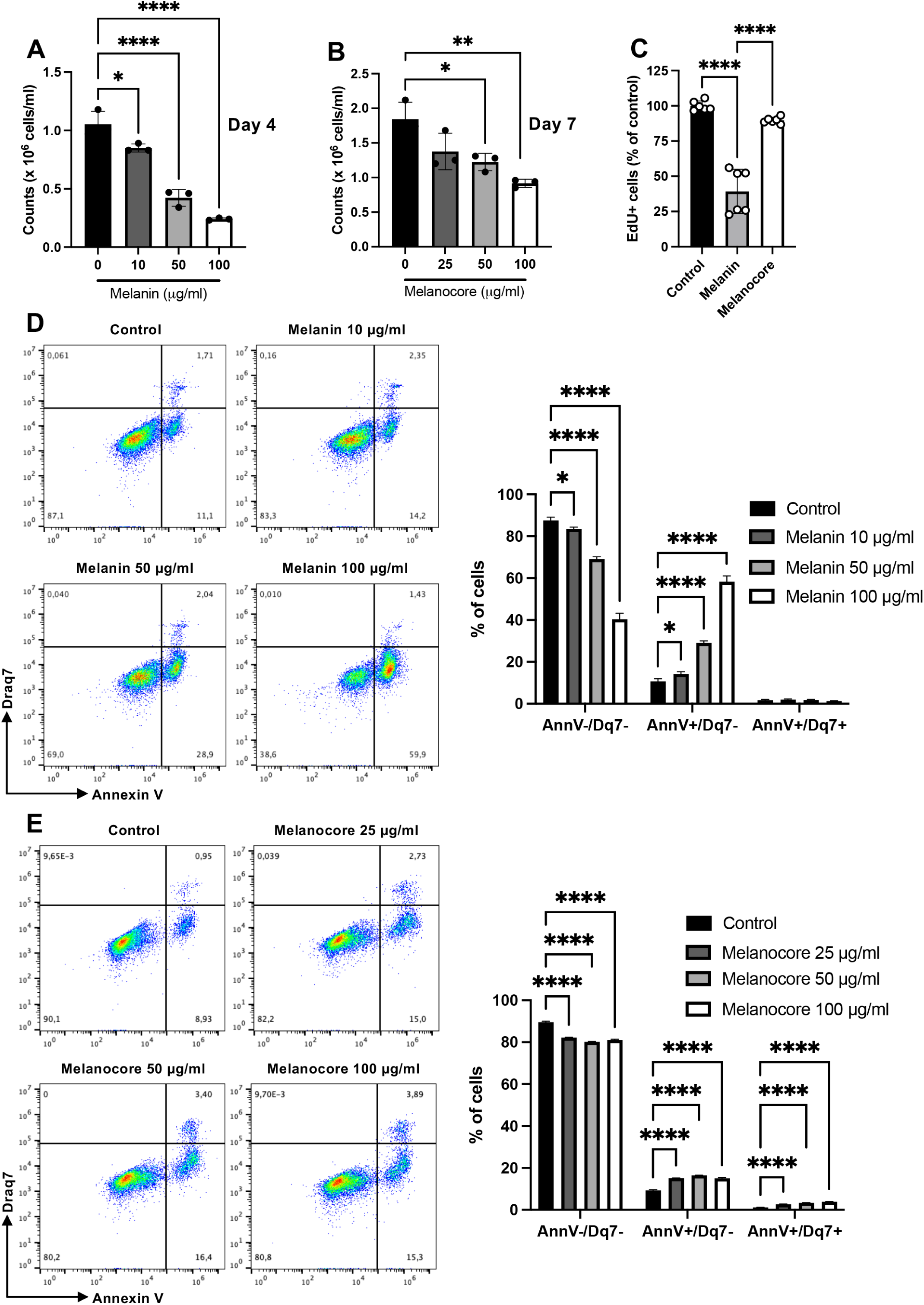
Melanin and melanocore-enriched conditioned medium impair MC growth through antiproliferative and pro-apoptotic mechanisms. (A) MCs were cultured with medium containing different concentrations of melanin. Cell counts were assessed after 4 days in culture. (B) MCs were cultured with medium containing different concentrations of melanocore-enriched medium (Melanocore) obtained from B16.F10 cells. Cell counts were assessed after 7 days in culture. (C) Cell proliferation assay based on EdU incorporation. (D-E) Cell viability assessment using Annexin V (AnnV) and Draq7 (Dq7) staining. Viable (AnnV^-^/Dq7^-^), apoptotic (AnnV^+^/Dq7^-^) and late apoptotic/necrotic (AnnV^+^/Dq7^+^) cells. The data are representative of at least three independent experiments and are given as mean values ± SEM (n=3). Unpaired t-test (A, B and C), two-way ANOVA (D and E); \**p*≤0.05; \*\**p*≤0.01; \*\*\**p*≤0.005; \*\*\*\**p*≤0.0001.

Melanin is secreted in the form of melanocores, which represent the membrane-free core of the melanosomes ^41, 42^. Hence, melanocores represent a physiologically relevant form of melanin in secretions, and we therefore investigated whether melanocores can affect MC growth. For this, we enriched melanocores from melanoma cell-conditioned medium and assessed whether the melanocore-enriched medium affected MC growth. Indeed, as seen in Fig. 2B, melanocore-enriched conditioned medium caused a dose-dependent inhibition of MC growth. Quantification of melanin using a fluorescence-based method showed that depletion of the conditioned media from melanocores caused a modest reduction of total melanin content, suggesting that melanin to a large extent is present in a soluble form. In agreement with this, it was noted that the melanocore-depleted conditioned media had only a slightly diminished capacity to inhibit MC proliferation (Suppl. Fig. 1).

We next asked whether the impact of free melanin and melanocores on MC growth is attributable to effects on proliferation and/or cell death, and this was assessed by EdU labeling and Annexin V/Draq7 staining, respectively. These assessments revealed that free melanin caused a significant reduction of MC proliferation, and a trend of impaired proliferation was also seen in response to the melanocore-enriched medium (Fig. 2C). It was also seen that free melanin had a significant pro-apoptotic impact on the MCs, as manifested by an increase in the Annexin V^+^/Draq7^-^ population and a corresponding decrease in viable (Annexin V^-^/Draq7^-^) cells (Fig. 2D). When assessing the effect of melanocore-enriched conditioned medium on cell viability, a slight but significant decrease in the proportion of viable cells was seen, accompanied by a corresponding increase in cells undergoing apoptosis-like cell death (Annexin V^+^/Draq7^-^) (Fig. 2E). Hence, these data suggest that melanin, both in its free form and in the form of melanocores has the ability to impair MC growth.

### Melanin and melanocores are taken up by MCs

To further outline how melanin affects MCs, we assessed whether melanin and/or melanocores can be taken up by MCs. For this, we first adopted the Fontana-Masson protocol to stain for melanin, and these analyses indicated that both melanin and melanocores are taken up by MCs (Fig. 3A). Melanin uptake by the MCs was also confirmed by quantification of cell-associated melanin, following the incubation of MCs with variable concentrations of melanin (Fig. 3B). Notably, melanin was found in the cytosol of the MCs, but we also noted the presence of melanin in the nucleus. To provide further insight into the latter, we used transmission electron microscopy (TEM). As seen in Fig. 3C-D, untreated MCs displayed the expected morphology, with the typical presence of large numbers of electron-dense secretory granules with dense core formation. After incubation of the MCs with melanin or melanocore-enriched conditioned medium, it was observed that the granule ultrastructure was affected, with an apparent loss of electron density. It was also noted that melanin could be found within granules, as well as in the cytosol. Further, the TEM analysis confirmed the presence of melanin/melanocores within the nucleus. After prolonged exposure (48 h) of the MCs to melanin or melanocores, we noted that melanin appeared to assemble into cytosolic protrusions directed towards the cell nucleus (Fig. 3D), and that melanin appeared to enter the nucleus via such protrusions through openings in the nuclear envelope (Fig. 3D).

**Fig. 3.**
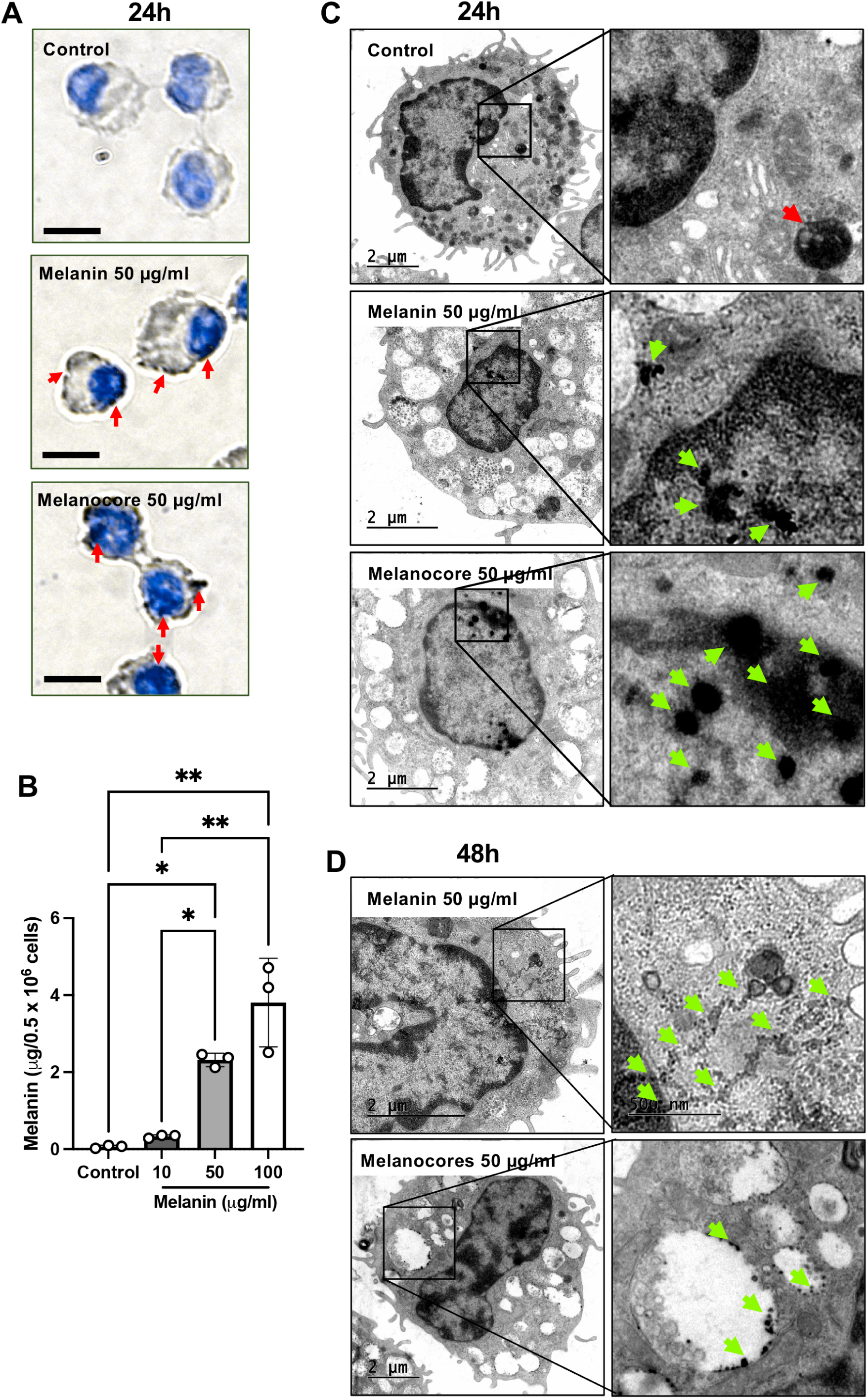
Melanin and melanocores are taken up by MCs. MCs were cultured under normal conditions or treated with medium containing 50 μg/ml of melanin or 50 μg/ml of melanocore-enriched medium (Melanocore). (A) Cytospin slides were stained with a combination of Fontana-Masson (black) and toluidine blue (blue). Cells were visualized using light microscopy. (B) MCs were cultured under normal conditions or treated with medium containing increasing concentrations of melanin; melanin concentration was measured in cell lysates (0.5 x 10^6^ cells) after 48 h incubation. (C) Transmission electron microscopy images showing the presence of dense core granule formation (red arrow) in control cells and the presence of melanin/melanocores (green arrows) in the cytoplasm, granules and nucleus of cells that had been treated with either melanin of melanocore-enriched conditioned medium (Melanocore). (D) Cytosolic protrusions directed towards the cell nucleus (green arrows) in cells treated with melanin or melanocore-enriched medium (Melanocore)for 48h. Note the presence of melanocores in granules.

To substantiate these finding we assessed if melanin is taken up by MCs also *in vivo*, in a melanoma context. To this end, we analyzed tumor tissue from mice in which subcutaneous melanoma tumors had been induced by inoculation of B16.F10 cells ^24^. Tissue sections from the mice were stained with Fontana-Masson and counterstained with toluidine blue. As expected, based on previous investigations (see Introduction), MCs could clearly be visualized at sites of tumor growth. The majority of the MCs were present in the tumor stroma, but MCs were also found within the tumor parenchyma (Fig. 4A-B). As expected, the tumor parenchyma stained intensely for melanin. When assessing for potential interactions between melanin and tumor-infiltrating MCs, we could not detect melanin in the vicinity of MCs located in the tumor stroma. In contrast, MCs located within the tumor parenchyma showed extensive interactions with melanin. Uptake of melanin into the tumor-infiltrating MCs was also observed (Fig. 4B and Suppl. Fig. 2). Hence, these findings suggest that melanin is taken up by MCs *in vivo*, in a melanoma tumor context.

**Fig. 4.**
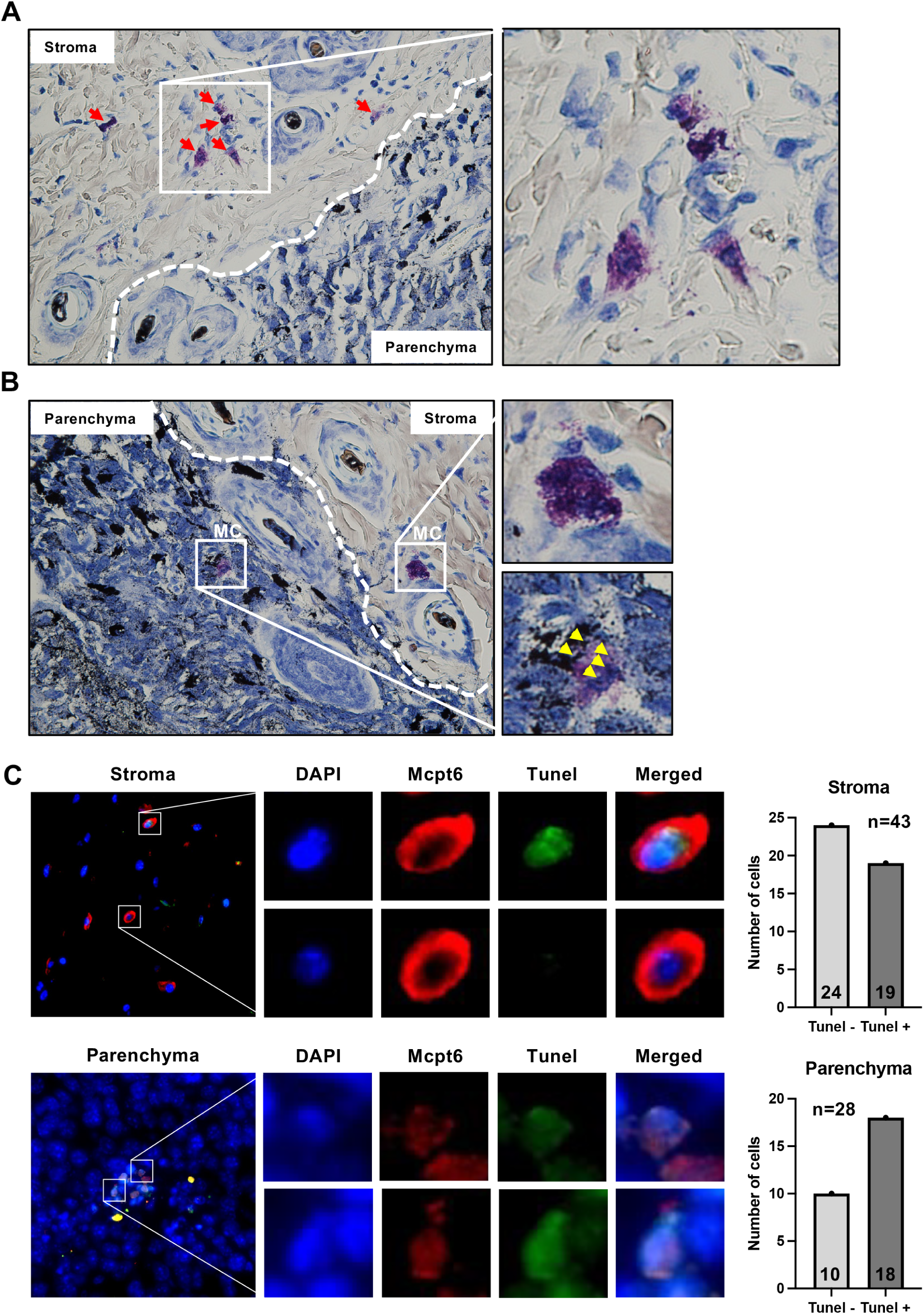
Melanin is detected within MCs present in the parenchyma of melanoma tumors. B16.F10 tumor samples were stained with a combination of Fontana-Masson (melanin – black) and toluidine blue (MCs – purple). Parenchyma and stroma areas are delimited by dashed white lines. (A) MCs were mostly found in the tumor stroma area (red arrows). (B) MC presence in the tumor parenchyma and stroma. Note the close contact between MCs and melanin (yellow arrows – lower left panel) in the parenchyma, but not in MCs present in the tumor stroma (lower right panel). (C) B16.F10 tumor samples were stained using a combination of *in situ* apoptosis detection (Tunel) and Mcpt6 immunofluorescence. Mcpt6-positive MCs were quantified and represented as either Tunel-negative (viable) or Tunel-positive (non-viable), as shown in the right panel.

To assess whether tumor-associated MCs were prone to undergo apoptosis, possibly as a result of melanin exposure, we stained MCs in both the tumor parenchyma and stroma for Tunel (an apoptosis marker). As shown in Fig. 4C, MCs located in the vicinity of melanoma cells (within the tumor parenchyma) were predominantly apoptotic (Tunel^+^), whereas MCs that were not in direct contact with melanoma cells (located in the tumor stoma) were largely Tunel-negative. Hence, these findings are compatible with a scenario in which melanin causes apoptosis of tumor-associated MCs.

### Effect of endocytosis inhibitors on melanin uptake by MCs

To address the mechanism of melanin uptake by MCs we assessed whether this could be mediated by endocytosis, which was evaluated by using endocytosis inhibitors. MitMAB, a specific dynamin inhibitor ^43^, had no cytotoxic effects on MCs in the absence of added melanin (Suppl. Fig. 3A). However, when MitMAB was added in the presence of melanin, the cytotoxic impact of melanin on MCs was markedly potentiated, as seen by a substantial decrease in the proportion of viable cells accompanied by a corresponding increase in the numbers of both Annexin V^+^/Draq7^-^ (apoptotic) and Annexin V^+^/Draq7^+^ (necrotic) cells (Suppl. Fig. 3A). In contrast, when assessing the effects of another dynamin inhibitor, Dyngo-4a ^44^, no potentiation of melanin-induced cytotoxicity could be observed (Suppl. Fig. 3B). To determine whether inhibition of endocytosis affects melanin uptake, untreated and MitMAB-treated cells were stained with Fontana-Masson and counterstained with DAPI. These analyses revealed a similar intensity of melanin staining in both the untreated and MitMAB-treated MCs (Suppl. Fig. 3C), suggesting that melanin is taken up by MCs independently of classical dynamin-mediated mechanisms. Previous studies have indicated that protease-activated receptor 2 (PAR-2) may act as an internalization receptor for melanin ^45^. To assess the impact of PAR-2 on melanin uptake in MCs we measured melanin uptake in the absence or presence of a PAR-2 antagonist (FSLLRY-NH_2_). However, melanin uptake was not affected by the PAR-2 antagonist (data not shown), suggesting that PAR-2 is not a major internalization receptor for melanin in MCs.

### Melanin affects the nuclear morphology in MCs and induces translocation of tryptase to the nucleus

Having shown that melanin can enter the nucleus of MCs, we next asked whether this can have an effect on nuclear features. For this purpose, we stained control-and melanin/melanocore-treated MCs with TO-PRO-3, a DNA-binding dye. These analyses revealed that both melanin and melanocore-enriched conditioned medium had profound effects on the intensity of TO-PRO-3 staining, with melanin causing a decrease whereas melanocores caused an increased staining intensity (Suppl. Fig. 4). It was also observed that both melanin and melanocores caused an increase in nuclear volume as compared with untreated cells (Suppl. Fig. 4).

In previous studies we have shown that MC tryptase, in addition to its presence within secretory granules, also can be found in the nucleus of MCs ^46, 47^. In agreement with these previous observations, tryptase (Mcpt6) was found in the MC nucleus at baseline conditions (in addition to its high abundance in granules)(Fig. 5A). Moreover, we have demonstrated that nuclear tryptase can have the capacity to regulate nuclear events ^46, 47^. Based on these findings, we hypothesized that the effects of melanin on nuclear events could be associated with translocation of tryptase to the MC nucleus. Indeed, as seen in Fig. 5A-B, treatment of MCs with B16.F10-conditioned medium caused a profound increase in the amount of nuclear tryptase. To verify this with an independent method, we performed a subcellular fractionation of MCs into cytosolic, membrane-bound (including granules), nuclear and chromatin-bound fractions. As expected, histone 3 (H3; a nuclear protein bound to chromatin), was highly abundant in the chromatin fraction but less prevalent in the other fractions (Fig. 5C). The different subcellular fractions were assessed for tryptase levels by immunoblotting. As seen in Fig. 5C, tryptase was found mainly in the cytosolic and membrane-bound (including granules) fractions under baseline conditions, but low levels of tryptase were also seen in the nuclear/chromatin fractions. However, after treatment of MCs with melanin or melanoma-conditioned medium, a marked increase in the levels of tryptase within the chromatin and nuclear fractions was observed, indicating translocation of tryptase into the nucleus (Fig. 5C). Next, we assessed whether this was selective for tryptase out of the multiple types of proteases that are present in the MC granules ^1, 48^. For this, we analyzed for CPA3, another highly abundant granule protease ^1, 48^. However, we did not note any melanin-induced translocation of CPA3 into the MC nucleus (Fig. 5C). Together, these findings show that melanin causes a selective translocation of tryptase into the MC nucleus.

**Fig. 5.**
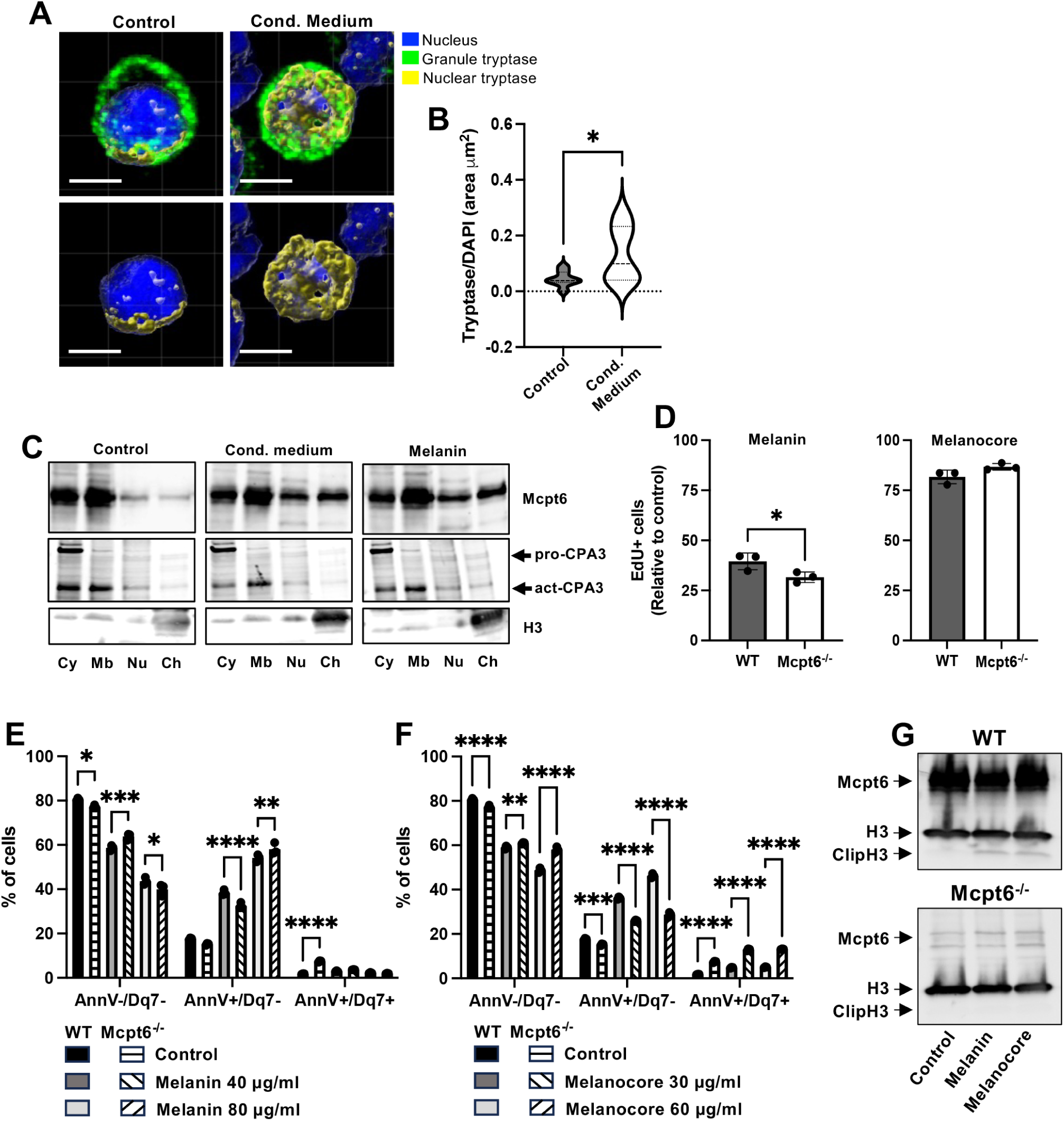
Melanin affects the nuclear morphology in MCs and induces translocation of tryptase to the nucleus. MCs were cultured either under normal conditions, in medium containing 30% B16.F10-conditioned medium or in medium containing either melanin or melanocore-enriched medium (Melanocores) at the indicated concentrations. (A) 3D images generated from confocal Z-stack sections stained for tryptase (green) and the nucleus (blue). The yellow color represents colocalization of tryptase and the nuclear dye (Hoechst 33342). Scale bars = 5 µm. (B) Violin plot showing the quantification of tryptase/nucleus colocalization area. (C) Western blot analysis of subcellular fractions: Cy – cytoplasmic proteins, Mb – membrane bound proteins, Nu – nuclear soluble proteins, Ch – chromatin bound proteins, for Mcpt6, CPA3 and histone 3 (H3). (D) Cell proliferation of WT vs. Mcpt6^-/-^ MCs, either under baseline conditions or after treatment for 48 h with 50 µg/ml melanin or melanocore-enriched medium (Melanocore). (E-F) Cell viability assessment using Annexin V (AnnV) and Draq7 (Dq7) staining. Viable (AnnV^-^/Dq7^-^), apoptotic (AnnV^+^/Dq7^-^) and late apoptotic/necrotic (AnnV^+^/Dq7^+^) cells. WT and Mcpt6^-/-^ cells were treated with different concentrations of melanin (E) or melanocore-enriched medium (F) for 8 days. (G) Effect of melanin and melanocore-enriched medium treatment on H3 clipping. Western blot analysis of WT and Mcpt6^-/-^ cells cultured under normal conditions or treated with medium containing 50 μg/ml of melanin (Melanin) or 50 μg/ml of melanocore-enriched medium (Melanocore). The presented data are representative of at least two/three independent experiments and are given as mean values ± SEM (n=3). \**p*≤0.05; \*\**p*≤0.01; \*\*\**p*≤0.005; \*\*\*\**p*≤0.0001. two-way ANOVA (E and F) and unpaired t-test (B and D).

We next assessed whether the absence of tryptase has an impact on the melanin-induced nuclear events. For this purpose, we developed MCs from both wild-type (WT) and tryptase-deficient (Mcpt6^-/-^) mice, and assessed their respective susceptibility to melanin/melanocores. These analyses demonstrated that the growth of Mcpt6^-/-^ MCs was less affected by melanoma cell-conditioned medium in comparison with WT cells (Suppl. Fig. 5A-C), which was reflected by less effects on EdU incorporation as compared with WT cells (Suppl. Fig. 5D). Moreover, in contrast to its effects on WT cells, melanoma cell-conditioned medium did not induce apoptosis in Mcpt6^-/-^ cells (Suppl. Fig. 5E). Notably though, melanin uptake was approximately equal in Mcpt6^-/-^ vs. WT cells (Suppl. Fig. 5F; compare with Fig. 3C). Further, the effect of melanin on nuclear staining intensity (TO-PRO-3) was modulated by the absence of tryptase, with blunted effects seen in Mcpt6^-/-^ vs. WT MCs (Suppl. Fig. 4). Likewise, the effect of melanocores on nuclear volume was less pronounced in Mcpt6^-/-^ vs. WT MCs, whereas free melanin had larger impact on nuclear volume in Mcpt6^-/-^ vs. control cells (Suppl. Fig. 4). Altogether, these findings suggest that the effects of melanin/melanocores on nuclear morphology are partly dependent on tryptase.

When assessing the impact of tryptase on effects induced by free melanin/melanocore-enriched conditioned medium, we noted that tryptase-deficient MCs were slightly but significantly more sensitive to melanin but marginally less sensitive to melanocores than were the WT cells (Fig. 5D). We also assessed whether the absence of tryptase had an impact on the ability of melanin/melanocores to induce cell death in MCs. As seen in Fig. 5E-F, melanin and melanocores induced similar extents of cell death in WT vs. tryptase-deficient MCs. However, it was notable that, whereas WT MCs predominantly underwent apoptosis-like cell death (Annexin V^+^/Draq7^-^) in response to melanin/melanocores, tryptase-deficient MCs mainly underwent necrosis-like cell death (Annexin V^+^/Draq7^+^)(Fig. 5E-F).

### Melanin induces tryptase-dependent histone 3 (H3) clipping in MCs

In previous studies we showed that tryptase present in the nucleus of MCs has the capacity to cause N-terminal H3 truncation (clipping) ^46, 47^. As seen in Fig. 5G, H3 clipping was virtually undetectable in MCs at baseline conditions. However, upon administration of either melanin or melanocores, markedly increased H3 clipping was noted (Fig. 5G). Considering the known ability of tryptase to execute H3 clipping, we asked whether the melanin/melanocore-induced H3 clipping was a consequence of translocation of tryptase to the nucleus. Indeed, when melanin of melanocore-enriched conditioned medium was incubated with tryptase-deficient MCs, H3 clipping was undetectable (Fig. 5G). Hence, melanin/melanocores induce tryptase-dependent H3 clipping in MCs.

### Melanoma-conditioned medium does not cause reduced expression of MC markers

Considering the marked impact of melanoma-conditioned medium/melanin on MC proliferation, we asked whether this can be associated with effects on the expression of MC marker genes. To this end, we assessed if B16.F10-conditioned medium can modulate the expression of Mcpt6 (tryptase), CPA3 or serglycin (a granule-localized proteoglycan). These analyses showed that B16.F10-conditoned medium did not suppress the expression of either the Mcpt6, CPA3 or serglycin genes (Fig. 6A). In fact, slightly increased expression of all these genes was noted in response to B16.F10-conditioned medium. Notably though, the levels of Mcpt6 and CPA3 protein were equal in the absence or presence of B16.F10-conditioned medium, indicating that the increase in mRNA expression for these genes in response to B16.F10-conditioned medium did not translate into effects at the protein level (Fig. 6B). Moreover, no significant effect on the expression of cell surface FcεRI (the high affinity IgE receptor) was seen after 4 days treatment of MCs with B16.F10-conditioned medium (Fig. 6C).

**Fig. 6.**
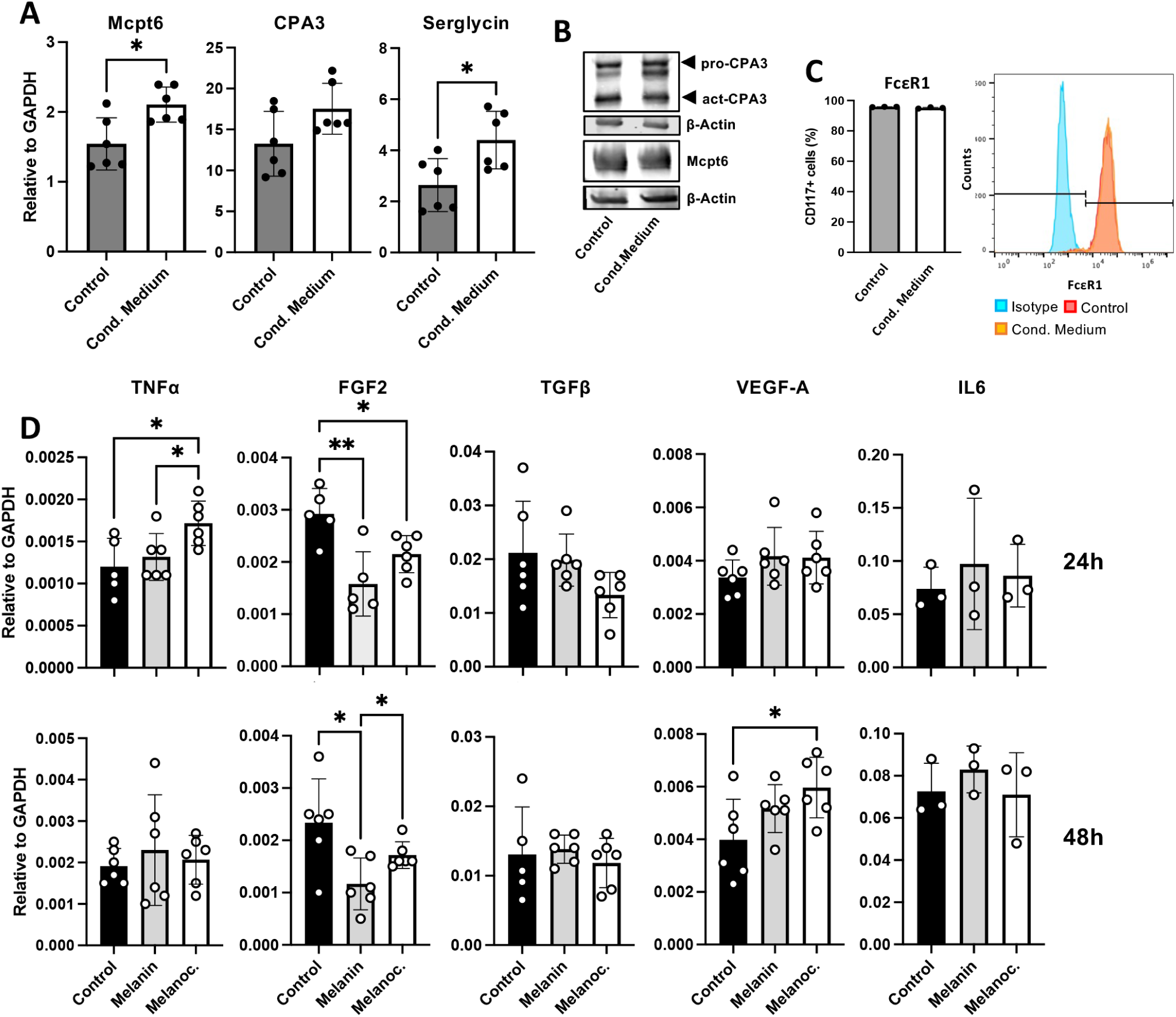
Effect of melanin on the expression of MC markers, cytokines and pro-angiogenic growth factors in MCs. MCs were cultured either under normal conditions, in medium containing 30% B16.F10-conditioned medium or in medium containing 50 μg/ml of melanin or melanocore-enriched medium (Melaonoc.). (A) Gene expression of the MC markers: tryptase (Mcpt6), CPA3 and serglycin, expressed as relative to the expression of GAPDH. (B) Western blot analysis of Mcpt6 and CPA3 in control cells and cells treated with 30% B16.F10-conditioned medium. (C) Cell surface expression of the high-affinity IgE membrane receptor (FcεR1) as determined by flow cytometry. (D) Expression of the tumor necrosis factor α (TNFα), fibroblast growth factor 2 (FGF2), transforming growth factor β (TGFβ), vascular endothelial growth factor A (VEGFa) and interleukin 6 (IL6) genes after 24 and 48 h treatment. The data are representative of at least three independent experiments and are given as mean values ± SEM (n=3). Unpaired t-test; \**p*≤0.05; \*\**p*≤0.01; \*\*\**p*≤0.005.

### Melanin and melanocores have differential effects on the expression of cytokines and pro-angiogenic growth factors in MCs

Next, we considered the possibility that melanin might affect the expression of cytokines and growth factors that potentially might support or repress tumor growth. Overall, these assessments revealed relatively modest effects of melanin on the expression of such genes in MCs (Fig. 6D). However, we noted that melanocore-enriched medium induced a slight but significant increase in the expression of the TNFα and VEGFa genes, and that both free melanin and melanocores caused a reduction in the expression of FGF2. In contrast, no significant effects of melanin/melanocores on the expression of either the TGFβ or IL-6 genes were observed (Fig. 6D).

### Melanin and melanocores suppress MC activation responses

In the following set of experiments, we asked whether melanin/melanocores, in addition to their negative impact on MC proliferation, also can modulate MC activation responses. To test this, we first analyzed whether melanin can affect the baseline expression of surface CD63, surface CD63 expression being an established marker for MC degranulation ^49^. These analyses showed that both melanin and melanocores caused a significant decrease in the baseline expression of CD63 on the MC surface, indicating suppression of spontaneous MC degranulation (Fig. 7A). Next, we performed experiments in which MCs were activated by either compound 48/80 (agonist for Mrgprb2; mouse ortholog to MRGPRX2 ^50^) or by IgE receptor crosslinking, followed by analysis of surface CD63 expression. As seen in Fig. 7B, CD63 surface expression was slightly enhanced following compound 48/80 administration, and both melanin and melanocores caused a suppression of the surface expression of CD63. In contrast, melanin or melanocores caused only minimal inhibition of surface CD63 expression in response to IgE receptor crosslinking (Fig. 7C).

**Fig. 7.**
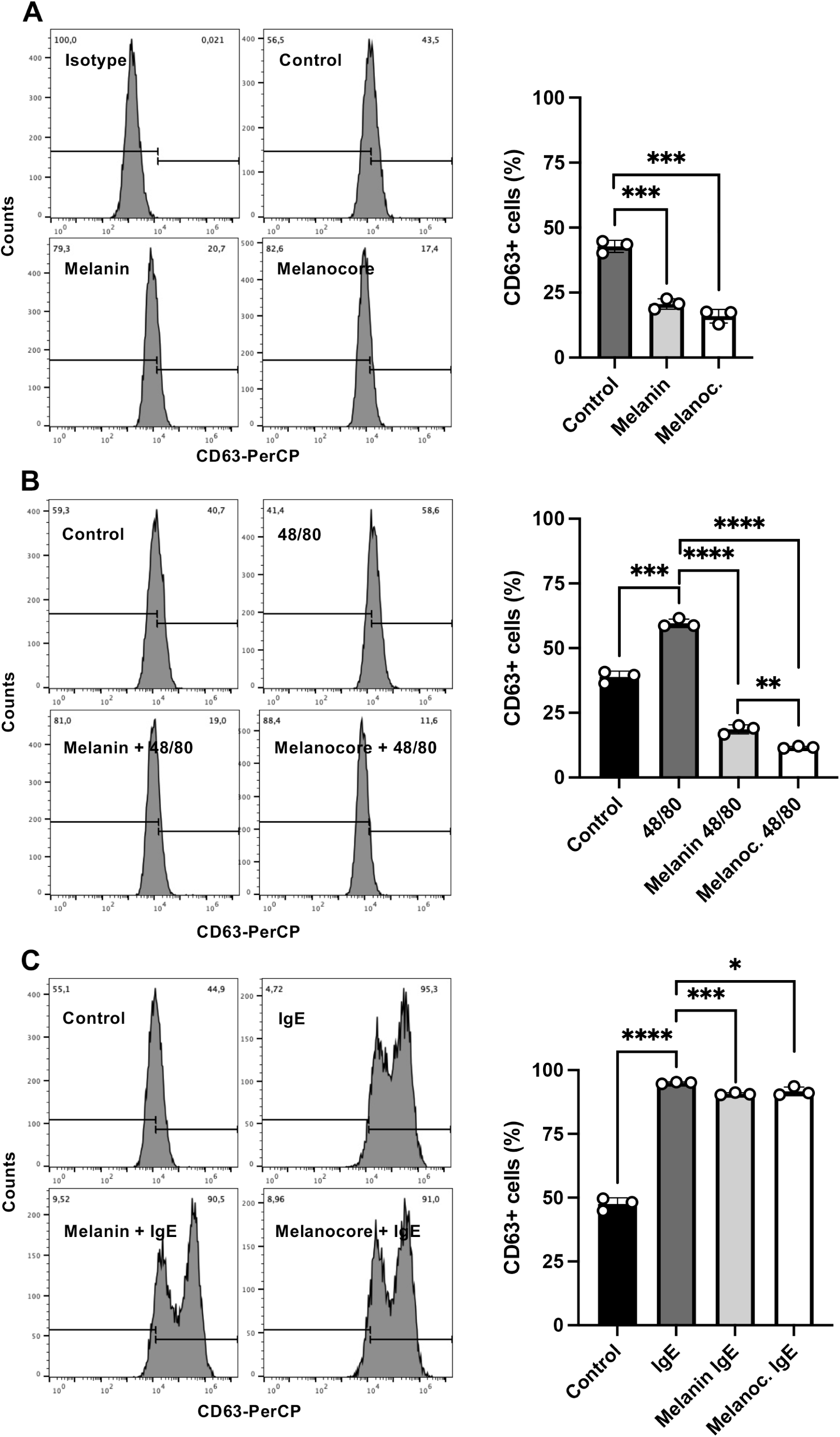
Effects of melanin and melanocores on MC activation responses. MCs were cultured under either normal conditions or in medium containing 50 μg/ml of melanin or melanocore-enriched medium (Melanoc). MC degranulation/activation was assessed by flow cytometry using an antibody against CD63. (A) Numbers of CD63^+^ MCs under normal and treatment conditions. (B) MCs under normal culture conditions or in the presence of melanin/melanocores were left untreated or treated with 50 μg/ml of compound 48/80 for 2 h. (C) MCs under normal culture conditions or in the presence of melanin/melanocores were sensitized with 0.1 μg/ml of IgE anti-DNP overnight, followed by challenge for 2 h with 0.5 μg/ml of DNP-HSA. The data are representative of at least two independent experiments and are given as mean values ± SEM (n=3). Unpaired t-test; \**p*≤0.05; \*\**p*≤0.01; \*\*\**p*≤0.005; \*\*\*\**p*≤0.0001.

Ligation of Mrgprb2/MRGPRX2 is emerging a major mechanism for MC activation in various contexts. Notably, Mrgprb2/MRGPRX2 expression is relatively low in bone marrow-derived MCs but is considerably higher in mature MCs of connective tissue subtype ^51^. To assess the effects of melanin on activation responses in MCs with high expression of Mrgprb2, we developed peritoneal cell-derived MCs (PCMC) and then assessed the effect of melanin/melanocore-enriched conditioned medium on their responses to activation via compound 48/80. Melanin/melanocores had only slight effects on their baseline CD63 expression (Suppl. Fig. 6A). In line with their higher expression of Mrgprb2 in comparison with bone marrow-derived MCs, the PCMCs responded more vividly to stimulation by compound 48/80 (Suppl. Fig. 6B). Moreover, it was seen that both melanoma and melanocore-enriched conditioned medium caused a strong suppression of CD63 expression in response to compound 48/80 stimulation (Suppl. Fig. 6B). Moreover, both melanin and melanocore-enriched conditioned medium strongly inhibited PCMC activation caused by IgE receptor crosslinking (Suppl. Fig. 6C), the latter thus in contrast to the bone marrow-derived MCs.

## Discussion

There is now a wealth of evidence supporting that MCs have the capacity to influence melanoma progression, either by supporting tumor growth or by having a protective function (see Introduction). In contrast to studies in which the impact of MCs on melanoma has been addressed, the possibility that melanoma cells may influence the function of MCs has to our knowledge not been investigated previously. Here we addressed this issue and demonstrate that the secretome of melanoma cells contains factors that can suppress MC growth, and our data suggest that melanin accounts for this effect. Potentially, this could represent a novel immunosuppressive function in melanoma, serving to reduce MC populations. This may thus add to previously known immunosuppressive mechanisms in melanoma settings, which include the recruitment of myeloid-derived suppressor-and T regulatory cells, impairment of MHC class I expression and secretion of immunomodulatory factors ^52, 53^.

Interestingly, several previous studies have shown an association between a high level of melanogenesis in melanoma and poor prognosis, and that melanogenesis can be associated with immunosuppression ^54, 55^. Based on the present study, we may thus speculate that high levels of melanin, as a consequence of highly active melanogenesis, can have a direct suppressive impact on tumor-associated MC populations. Melanin-induced MC suppression may thus account, at least partly, for the negative impact of melanogenesis on disease outcome.

Most likely, a suppressive impact of melanin on MCs would be most prominent in the close vicinity to the melanoma cells, i.e. within the tumor parenchyma. In line with this, we noted that MCs within the parenchyma of subcutaneous melanoma tumors engage in close contacts with melanin, whereas MCs found in the tumor stroma display less contacts with melanin.

Notably, MCs are markedly more abundant in the tumor stroma than within the tumor parenchyma ^16, 19, 24^ and, on a speculative angle, we may propose that the low numbers of MCs within the tumor parenchyma may, at least partly, be linked to suppressive effects of melanin.

The mechanism by which melanin suppresses MC growth is intriguing. Our results suggest that melanin, either in its free form or in the form of melanocores, has a profound inhibitory impact on MC proliferation, and also that melanin/melanocores induce MC apoptosis. In the skin, melanin secreted by melanocytes is taken up by keratinocytes ^56^ and, based on this notion, we asked whether melanin also can be taken up by MCs. Indeed, our findings reveal that melanin is vividly taken up by MCs, as seen both in cultured MCs and *in vivo* in subcutaneous B16.F10 melanoma tumors. Further, it was seen that melanin, after uptake, can enter the MC nucleus, and it was also noted that the nuclear morphology was profoundly affected by the uptake of melanin. We may thus envision that the antiproliferative effect of melanin on MCs may be related to effects on the nuclear compartment.

To pinpoint the mechanism of uptake we considered the possibility that melanin was taken up by dynamin-dependent pathways such as endocytosis. However, dynamin inhibition did not intervene with melanin uptake, suggesting a dynamin-independent mechanism ^57^. Clearly, extended investigations will be warranted to fully clarify the exact mechanism of melanin uptake into MCs. With regard to the mechanism of melanin uptake into keratinocytes, it has been suggested that melanin, in the form of melanocores, may be taken up by phagocytosis ^45^. However, since phagocytosis is generally considered to be dynamin-dependent, it is difficult to reconcile our present findings with a mechanism where melanin is taken up by MCs via conventional phagocytosis. Interestingly, previous investigations have suggested a role for filopodia-associated E-cadherin in the transfer of melanin into keratinocytes ^58, 59^. We cannot exclude that a similar mechanism is operative in melanin uptake into MCs. However, MCs would not normally express E-cadherin, and such a mechanism of melanin uptake thus appears less likely.

In previous investigations we have shown that MC tryptase, in addition to its location within the secretory granules, also can be found in the nucleus and can regulate cell proliferation ^46,47^. Based on these findings, we hypothesized that the effects of melanin on MCs possibly could be linked to the presence of tryptase within the nuclear compartment. Indeed, our findings reveal that melanin causes an extensive translocation of tryptase into the MC nucleus. Notably, this translocation appeared to be selective for tryptase among the various proteases that are abundant within the MC secretory granules. To determine whether tryptase can account for the melanin-mediated effects on MCs, we assessed whether melanin had a differential impact on WT vs. tryptase-deficient MCs. These analyses revealed that tryptase-deficient MCs were less sensitive to the growth inhibitory action of melanin as compared with WT cells, and also that the absence of tryptase resulted in a shift from apoptosis-like-to necrosis-like cell death in response to melanin, the latter being in line with previous findings^60^. Further, the effects of melanin on the nuclear morphology in MCs was partly dependent on tryptase. Hence, these findings indicate that tryptase partly accounts for the effects of melanin on MCs. Clearly, these findings are in line with a previous report showing that melanoma growth in vivo is affected by the absence of tryptase ^24^.

In previous studies we have shown that tryptase located in the MC nucleus has the capacity to cause N-terminal clipping of H3 and other core histones ^46, 47^. In agreement with this, H3 clipping was seen under baseline conditions in this study and, interestingly, increased H3 clipping was observed in response to melanin. Further, we demonstrate that the H3 clipping was critically dependent on MC tryptase. The N-terminal ends of H3 (and other core histones) are important sites for modification by epigenetic marks, including acetylation, methylation and phosphorylation ^61, 62^. Hence, a likely scenario would be that increased levels of nuclear tryptase in response to melanin can result in the eradication of selected epigenetic histone marks, and it is possible that such events may, at least partly, contribute to the molecular mechanism(s) by which melanin reduces MC growth.

As an additional level of MC suppression, we show that melanin reduces the ability of MCs, both bone marrow-derived MCs and PCMCs, to undergo activation in response to Mrgprb2 stimulation. In contrast, melanin did not affect the responsiveness of bone marrow-derived MCs to IgE-mediated activation, whereas robust inhibition of IgE-mediated responses was seen in MCs of PCMC type. The latter is thus in agreement with a previous study in which melanin was shown to impair the response of a MC-like cell line to activation via IgE receptor crosslinking ^63^. It is presently not known by which mechanism(s) MCs are activated within melanoma tumors *in vivo*. Notably though, Mrgprb2 is a major cell surface receptor expressed by MCs, and it is known that Mrgprb2 responds to multiple types of stimulants ^50, 51^. It thus appears likely that Mrgprb2 might play an important physiological role in MC activation responses in melanoma settings, and its inhibition by melanin might consequently suppress the ability of MCs to undergo activation.

Altogether, our findings reveal that melanin has the capacity to suppress MC populations both at the level of growth and activation. Potentially, these effects of melanin could serve to minimize the ability of MCs to influence melanoma progression.

## Acknowledgements

This study was supported by grants from The Swedish Research Council, The Swedish Cancer Foundation, The Swedish Heart and Lung Foundation, The Erling-Persson Foundation and The Knut and Alice Wallenberg Foundation.

## Author Contributions

FRM conceived of and designed the study, performed most of the experimental work, interpreted data and contributed to the writing of the paper; LN performed experiments; IÖM performed experiments; MG performed experiments; GP contributed to the design of the study, interpreted data and wrote the manuscript.

## Legends to Supplemental Figures

**Suppl. Fig. 1. Effect of melanocore-depleted B16.F10-conditioned medium on MC proliferation.** (A) Melanin concentration was measured in B16.F10-conditioned medium, before and after melanocore depletion. (B) Cell proliferation assay based on EdU incorporation in cells cultured for 48 h under normal conditions, in medium containing 30% B16.F10-conditioned medium, or in 30% melanocore-depleted B16.F10-conditioned medium. Unpaired t-test; \**p*≤0.05; \*\**p*≤0.01; \*\*\**p*≤0.005; \*\*\*\**p*≤0.0001.

**Suppl. Fig. 2. MCs present in the parenchyma of melanoma tumors are closely associated with melanin.** B16.F10 tumor sections were stained with a combination of Fontana-Masson (melanin – black) and toluidine blue (MCs – purple). Note that MCs present in the parenchymal engage in close contacts with melanin (yellow arrows).

**Suppl. Fig. 3. Effects of endocytosis inhibitors on melanin uptake by MCs.** MCs were pre-treated for 1 h with the endocytosis inhibitors MitMAB (3 µM; A) or Dyngo-4a (10 µM; B) followed by treatment with medium containing 50 μg/ml of melanin for 24 h. Cell viability was assessed by Annexin V (AnnV) and Draq7 (Dq7) staining. (C) Cytospin slides showing untreated controls and cells treated with medium containing 50 μg/ml of melanin in the absence or presence of 3 μM MitMAB. Cytospin slides were stained with Fontana-Masson (black) and DAPI (blue) and examined using a combination of fluorescence and DIC bright field microscopy. The data are representative of at least three independent experiments and are given as mean values ± SEM (n=3). Two-way ANOVA; \**p*≤0.05; \*\**p*≤0.01; \*\*\**p*≤0.005; \*\*\*\**p*≤0.0001.

**Suppl. Fig. 4. Melanin affects the nuclear morphology in MCs.** Staining of control- and melanin/melanocore (50 µg/ml)-treated MCs (WT or Mcpt6^-/-^) with the TO-PRO-3 nuclear dye. (A) Z-stacks of TO-PRO-3-stained cells were used to generate 3D images of cell nuclei after 24h treatment with melanin or melanocore-enriched medium. (B; left panel) Mean fluorescence intensity per nucleus. (B; right panel) Nuclear volume in μm^3^. Two-way ANOVA; \**p*≤0.05; \*\**p*≤0.01; \*\*\**p*≤0.005; \*\*\*\**p*≤0.0001.

**Suppl. Fig. 5. Effect of melanin on the growth of tryptase-deficient MCs.** Tryptase-deficient (Mcpt6^-/-^) MCs were cultured under normal conditions (Control) or in medium containing 30% B16.F10-conditioned medium (Cond. Medium). (A) MCs were cultured for 7 days, and cell counts were performed at day 0 and after 7 days in culture (right panel). (B) MC cultures were followed for 18 days. Medium was replaced every 3^rd^/4^th^ day, cells were counted and diluted to 0.5x10^6^ cells/ml after each passage. (C) Cell size in μm. (D) Cell proliferation assay based on EdU incorporation. (E) Cell viability assessment using Annexin V (AnnV) and Draq7 (Dq7) staining. Viable cells (AnnV^-^/Dq7^-^), apoptotic cells (AnnV^+^/Dq7^-^) and late apoptotic/necrotic cells (AnnV^+^/Dq7^+^). (F) Melanin concentration measured in lysates (0.5 x 10^6^ cells) of cells treated with medium containing increasing concentrations of melanin for 48 h. The data are representative of at least two/three independent experiments and are given as mean values ± SEM (n=3). Unpaired t-test; \**p*≤0.05; \*\**p*≤0.01; \*\*\**p*≤0.005; \*\*\*\**p*≤0.0001.

**Suppl. Fig. 6. Effect of melanin and melanocores on activation responses in peritoneal cell-derived MCs (PCMCs).** PCMCs were cultured under either normal conditions or in medium containing 50 μg/ml of melanin or melanocore-enriched conditioned medium (Melanocore). PCMC degranulation/activation was assessed by flow cytometry using an antibody against CD63. (A) Numbers of CD63^+^ PCMCs under normal and treatment conditions. (B) PCMCs under normal culture conditions or in the presence of melanin/melanocores were left untreated or treated with 50 μg/ml of compound 48/80 for 2 h. (C) PCMCs under normal culture conditions or in the presence of melanin/melanocores were sensitized with 0.1 μg/ml of IgE anti-DNP overnight, followed by challenge for 2 h with 0.5 μg/ml of DNP-HSA. The data are representative of at least two independent experiments and are given as mean values ± SEM (n=3). Unpaired t-test; \**p*≤0.05; \*\**p*≤0.01; \*\*\**p*≤0.005; \*\*\*\**p*≤0.0001.

